# A high fat containing chicken egg-only diet suppresses fatty liver induced by a lipid-rich methionine and choline deficient diet

**DOI:** 10.1101/2020.06.14.151316

**Authors:** Ken-ichi Isobe, Naomi Nishio, Ami Kuzuya, Kana Kato, Aki Hatanaka, Rena Suzuki, Miki Kawai, Aoi Kanjya, Chiaki Suitou, Yui Nakano, Manae Nagasaki

## Abstract

Nonalcoholic fatty liver disease (NAFLD) is a simple hepatic steatosis, which may proceed to nonalcoholic steatohepatitis (NASH) in the presence of steatosis and inflammation with hepatocyte injury. Obesity induced by a high-fat diet or high monosaccharide diet is considered a risk factor for NAFLD. A popular mouse model of NAFLD is a high-fat diet consisting of 60% energy from fat. A modified methionine and choline deficient (MCD) diet containing 60% energy from fat (CDAHFD) is a quick induction model for NAFLD. Chicken eggs also contain 60% energy from fat with very low carbohydrate. Here we compared liver pathologies in mice fed either a CDAHFD or egg-only diet. We found that a CDAHFD induced NAFLD within only two weeks.

Ballooning of hepatocytes with an of immune cell appearance in the liver and high serum ALT and AST indicated that the mice fed CDAHFD developed NAFLD, which could proceed to NASH. However, mice fed an egg-only diet did not develop NAFLD even after 7 weeks. These mice showed normal liver histology with normal ALT and AST. The mice fed an egg-only diet showed high blood ketone bodies and normal blood glucose. Furthermore we found that the mice fed a combination CDAHFD /egg diet or mice fed an egg-only diet after two weeks of CDAHFD diet showed almost normal ALT and AST with reduced levels of fat bodies in the liver. These results indicate that an egg-only diet strongly inhibits high fat and carbohydrates induced NAFLD.

## Introduction

Obesity and metabolic syndrome represent serious worldwide problems. Type 2 diabetes, nonalcoholic fatty liver disease (NAFLD), atherosclerosis, stroke, and ischemic heart disease occur in people with these conditions (1). NAFLD is a simple hepatic steatosis, which may proceed to nonalcoholic steatohepatitis (NASH) in the presence of steatosis and inflammation with hepatocyte injury (ballooning) (2, 3). NASH can progresses to liver cirrhosis and hepatocellular carcinoma (HCC).

One characteristics of patients with nonalcoholic steatohepatitis is an increased tendency to develop hepatocellular carcinoma (4). In developed counties including USA, Japan and those of Europe, more than one fourth of the population has been estimated to be affected by NAFLD (5, 6). Although high calorie consumption induces NAFLD in modernized countries, energy sources and other nutritional factors may also affect disease outcomes (7). To study these conditions several murine models have been developed to investigate the mechanism of NALFD and NASH.

In vivo studies identified that long-term exposure to a high-fat diet (HFD) led to NASH as well as a high rate of spontaneous hepatocellular carcinomas in C57BL/6J mice but not in A/J mice (8). In contrast the NASH phenotype induced by methionine and choline deficient (MCD) diet is more severe in A/J mice than in C57BL/6 mice (9). The MCD diet model is one of the most common tools for NAFLD/NASH research (10). Originally the MCD diet contained about 40% sucrose and 10% fat. This diet induced the rapid onset of liver cell death and oxidative stress as well as the production of cytokines leading to NASH (11, 12, 13).

However, this dietary methionine-/choline-deficient mouse model can cause severe weight loss (14, 15), which is not a characteristic of NASH seen in human patients. To overcome weight loss of MCD diet, a new lipid-rich MCD model was developed a choline-deficient, L-amino acid-defined, high-fat diet (CDAHFD). This diet is high in fat (60 kcal% fat) with limited amounts of methionine (0.1%) and a lacks of choline (16).

Interestingly chicken eggs also contain high amounts of lipid (60 kcal%) with a very low amount of carbohydrates. Here we compared the development of NAFLD in both a CDAHFD and an egg-only diet. We observed that although the mice fed a CDAHFD showed marked fat deposition in the liver, the mice fed eggs only showed no pathological changes in their livers either serologically or histologically. In addition, we also examined whether an egg-only diet can reverse fatty liver caused by a CDAHFD.

## Materials and Methods

### Mice

Eight-week-old C57BL/6 (C57BL/6n; B6) male mice were purchased from SLC Japan. They were maintained on either a normal diet (ND; CLEA Rodent Diet CE-2) or on an egg-only diet, under 12-hour light and dark cycles and specific pathogen-free conditions in the Animal Research Facility at the Nagoya Women’s University, and were used according to institutional guidelines. All mice were housed with up to 5 animals per cage, with ad libitum access to diet and tap water. The Animal Care and Use Committee of Nagoya Women’s University approved the study protocol.

### Mouse diets and feeding

The CLEA Rodent Diet CE-2 was as the ND; it contains 8.84% water, 25.48% protein, 4.61% fat, and 61.07% carbohydrates, along with vitamins and minerals. The energy from 100g of CE-2 is 339.1 kcal (Chubu Kagaku, Japan). The CDAHFD diet (A06071302) is L-amino acid diet with 60 kcal% fat with 0.1% methionine and no added choline (Research Diets, New Brunswick USA). For the egg-only diet, we used boiled chicken eggs. Eggs were obtained from firms in Aichi, Japan. Nutritional analysis of the boiled eggs was performed at Japan Food Research Laboratories (Nagoya, Japan). Boiled eggs have a high water content (75.6g/100g), and 12.9g/100g protein, 9.6g/100g fat and 1.1g/100g carbohydrate in addition to vitamins and minerals; 36.1% of calories (E) from protein, 60.8% E from lipid and 3.1% E from carbohydrates.

### Histological analysis

Liver tissue was collected and fixed in 4% paraformaldehyde buffered with PBS (pH 7.4) and blocked in paraffin. Paraffin-embedded 4-μm sections were stained with hematoxylin for 5minutes and with eosin for 3minutes.

### Biochemical analysis

Serum samples were obtained from the mice at the time of sacrifice. Serum glucose, ketone bodies, and the liver marker enzymes alanine transaminase (ALT) and aspartate transaminase (AST) were measured by enzymatic colorimetric assays. Serum levels of ALT and AST were calculated by measuring pyruvate (555nm) using WAKO assay kits (Wako Pure Chemicals, Osaka, Japan). Blood glucose was measured by using the LaboAssay™ Glucose kit (Wako Pure Chemicals, Osaka, Japan). Ketone bodies, including total ketone bodies (T-KB) and 3-hydroxybutyrate (3-HB) were measured with the Wako Autokit. Total Ketone Bodies and 3-HB (Wako Pure Chemicals, Osaka, Japan). Total cholesterol, HDL cholesterol and triglycerides (TG) were measured using the Wako E test, and free fatty acids (FFA) and non-esterified fatty acids (NEFA) were measured using the NEFA C test (Wako Pure Chemicals, Osaka, Japan). Wavelengths were measured with an xMark™ microplate absorbance spectrophotometer (Bio Rad).

### Statistical analyses

Data are expressed as the means□ ±□ standard deviations (SD). Statistical comparisons were performed using a Student t-test or a one-way analysis of variance (ANOVA) and Tukey-Kramer test. P-values<0.05 were considered statistically significant.

## Results

### Sixty percent energy from fat in the CDAHFD induces NAFLD, but the same amount of fat from eggs does not

We compared the inducibility of NAFLD by a high-fat diet between the CDAHFD and egg-only diet. The differences between the two included the low carbohydrates in the egg-only diet and the low methionine and lack of choline in the CDAHFD. Both diets included around 60 % energy from fat. Eight-week-old male mice were raised on an CDAHFD or egg-only diet for 7 weeks. After 4 weeks some mice were sacrificed for histological examination of the liver. Other mice were sacrificed at 7 weeks for examination of organ appearance, organ histology, and metabolic analysis (Figure 1a). The body weights of the mice fed CDAHFD and those of the mice fed only eggs were both increased, although the body weight increase of the CDAHFD mice was slightly lower than that of the mice fed an egg-only diet (Fig. 1b). This difference in body weights might be caused by the amount of food consumed as the mice were fed ad libitum and the amount of eggs consumed per mouse was approximately twice that of the CDAHFD (data not shown). The mice fed a CDAHFD appeared slightly smaller than the mice fed only eggs after 7 weeks (Fig.1c). The livers of the mice fed eggs only looked bigger than those of the mice fed a CDAHFD for 7 weeks. However, the weights of other organs were almost the same (Fig.1d). Liver histology showed that in mice fed a CDAHFD, liver cells contained big and small vacuoles after only 4 weeks, whereas the liver cells of egg-only mice appeared almost normal at both 4 weeks and 7 weeks (Fig.1e). Moreover serum ALT and AST in the CDAHFD mice were higher than those of the egg-only mice (Fig.1f).

**Figure 1.**
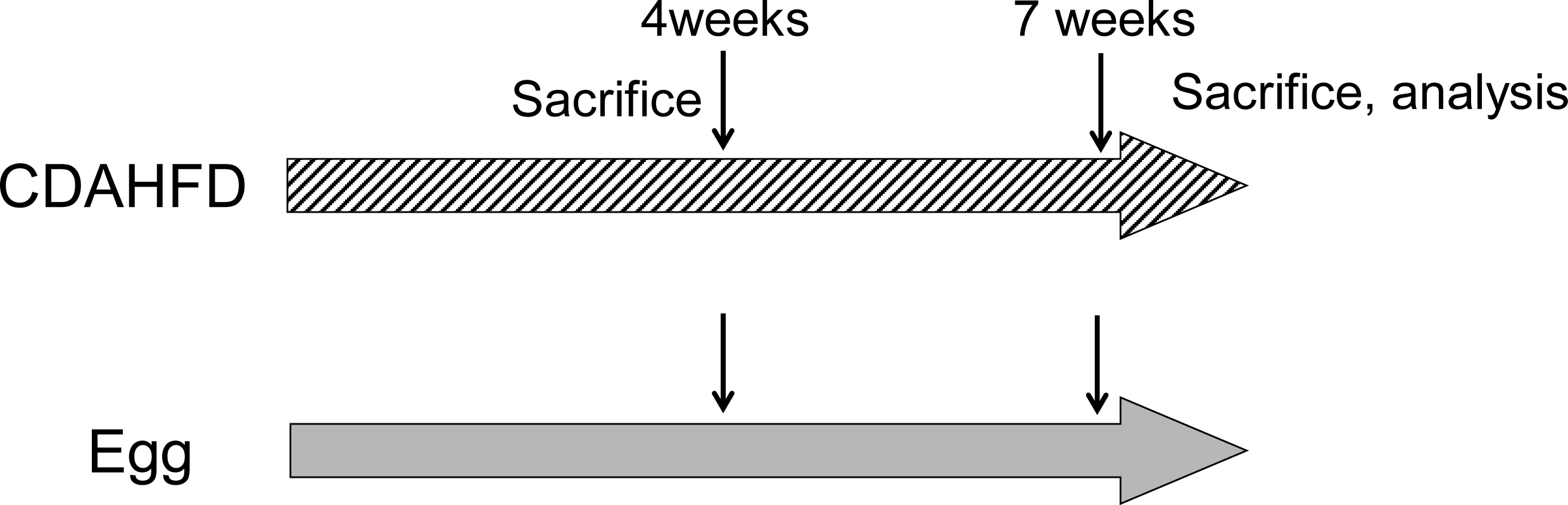
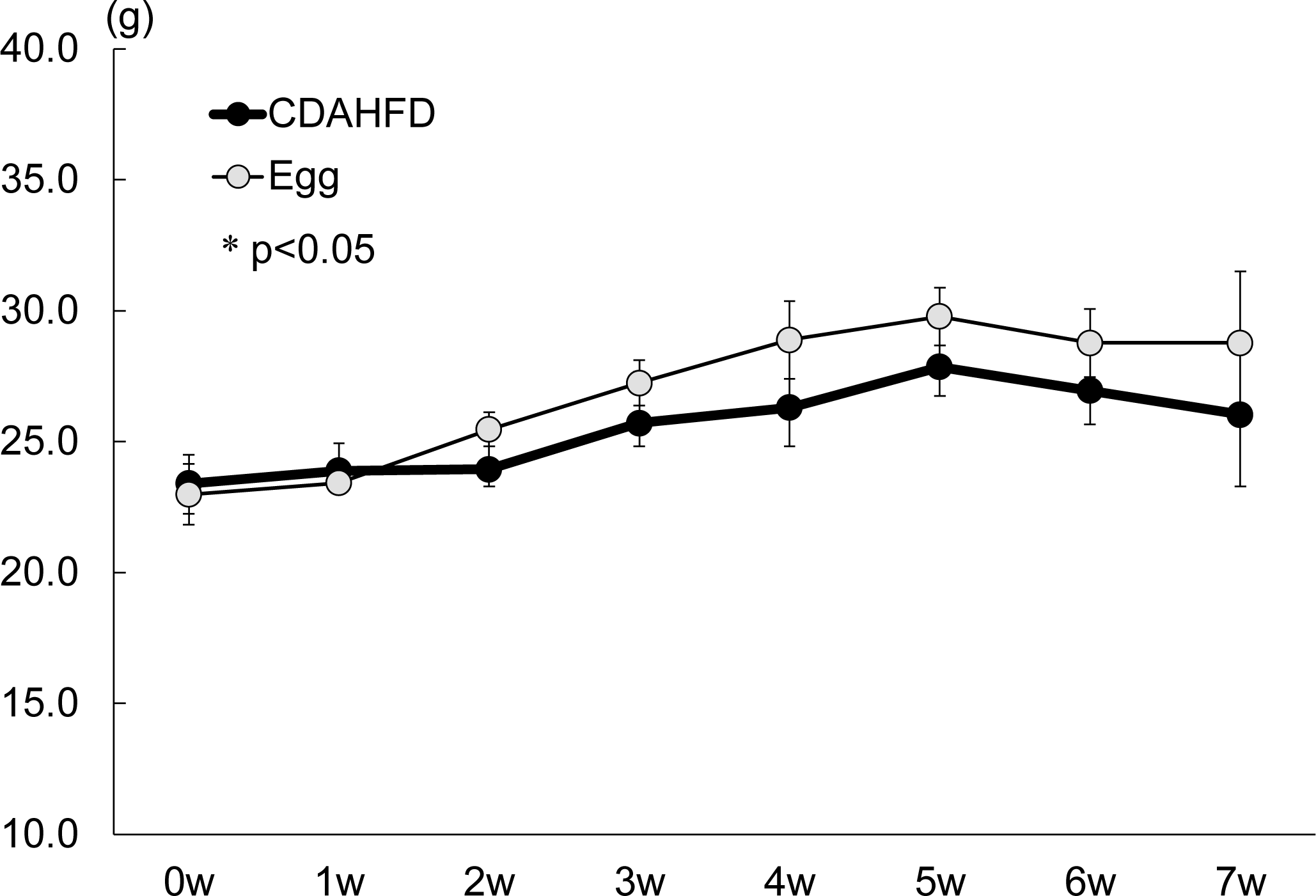
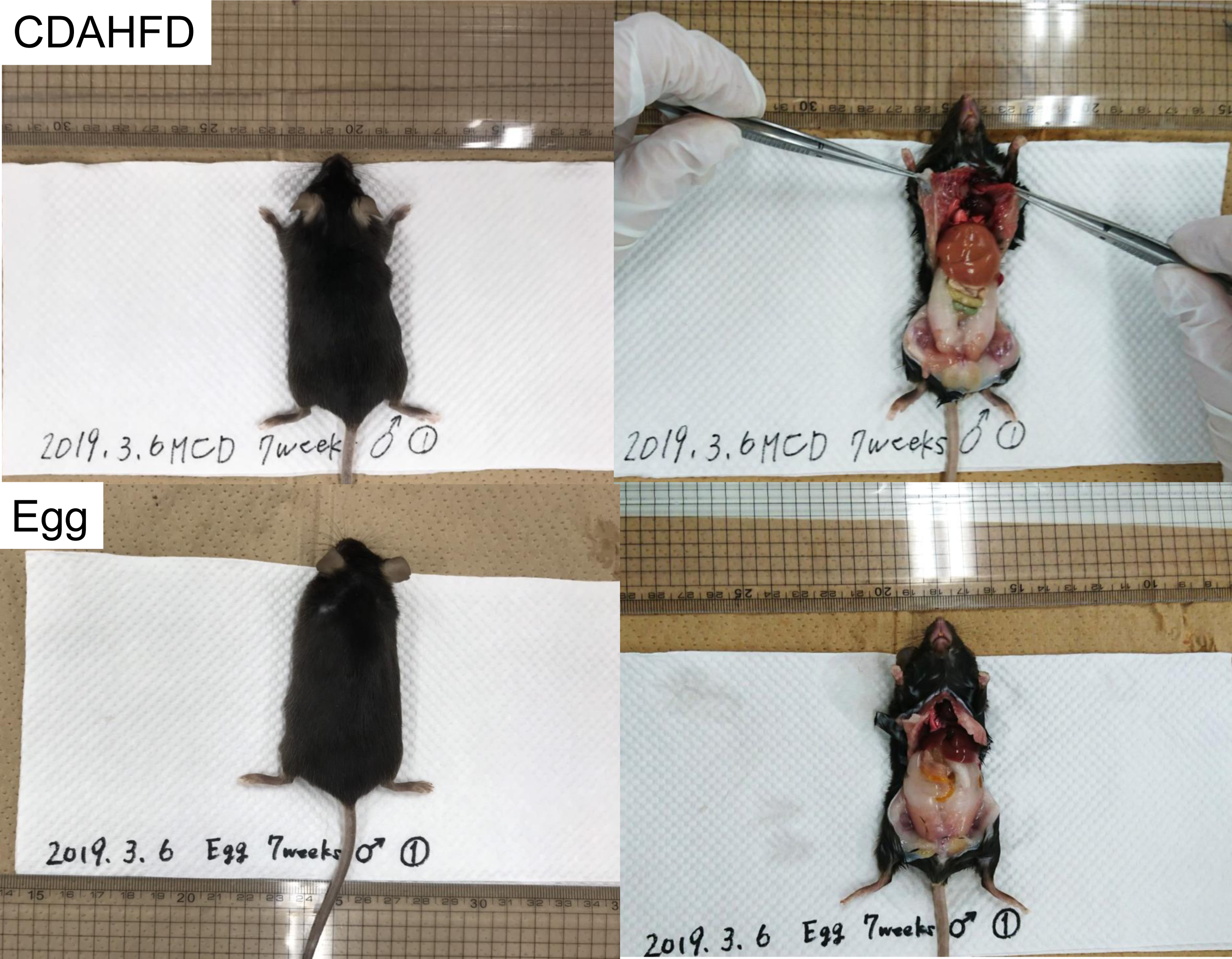
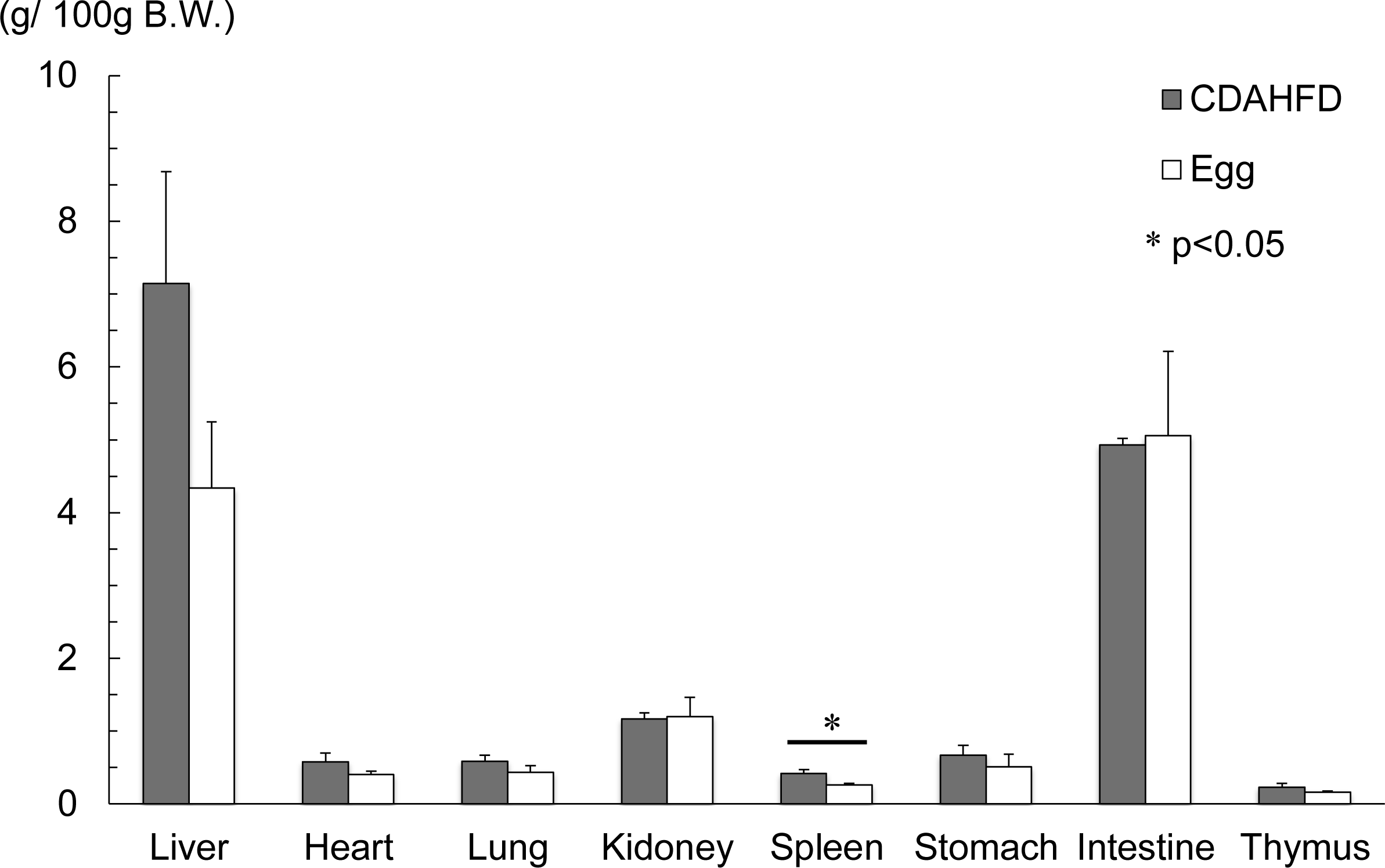
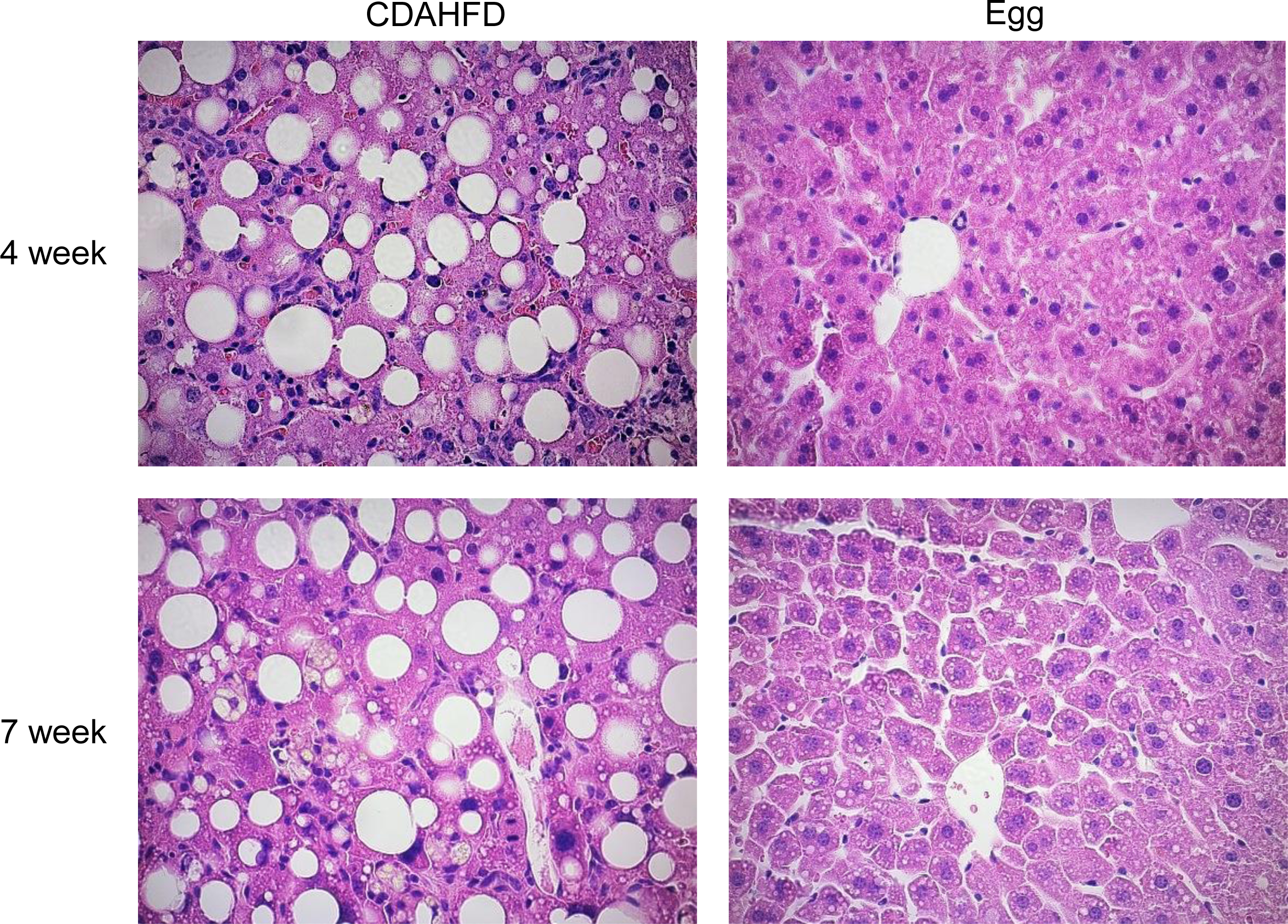
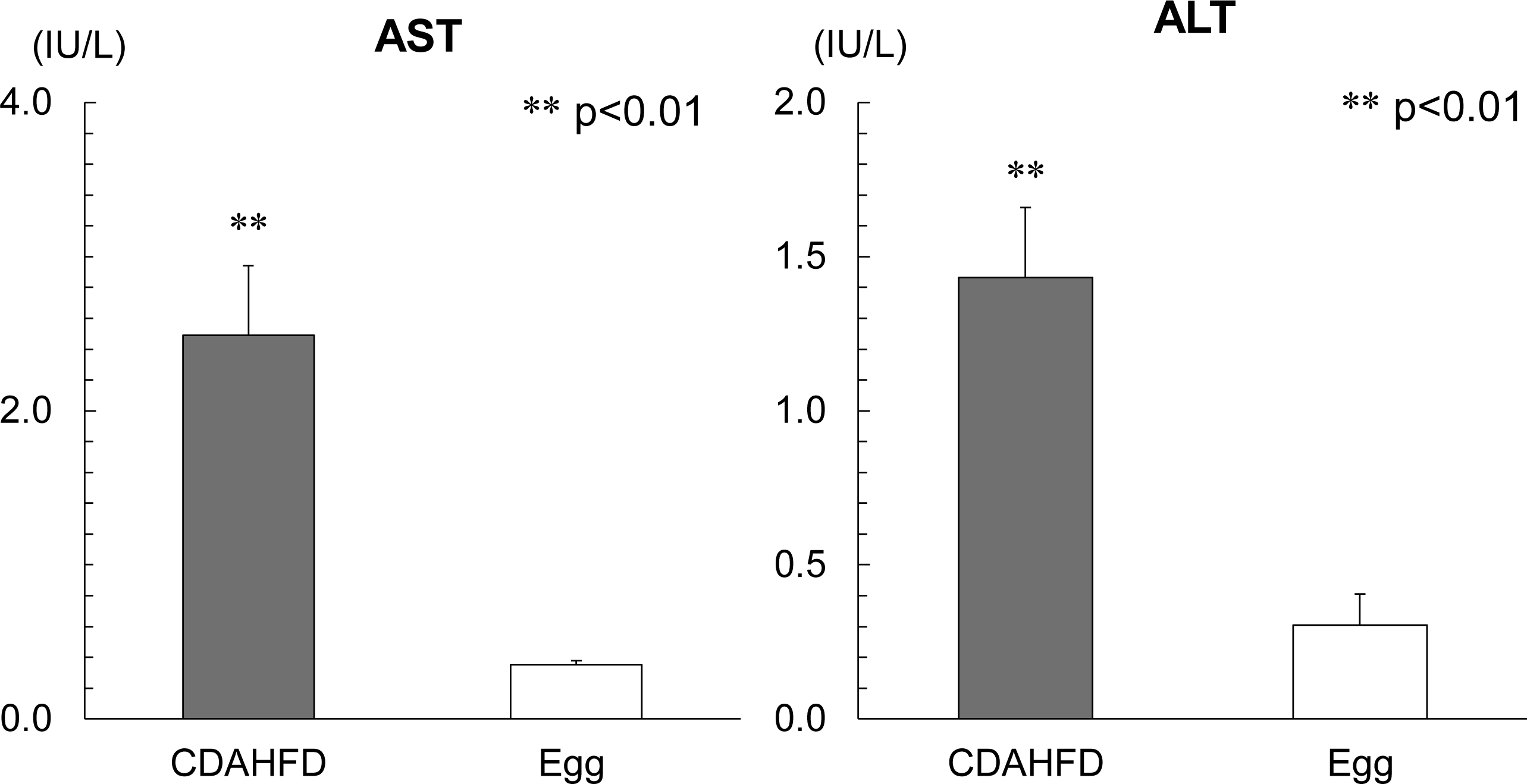
NAFLD was induced in mice fed a CDAHFD but not in egg-only mice. (a) Eight-week-old male mice were fed either CDAHFD or eggs only ad libitum for 7 weeks. (b) Body weights of CDAHFD or egg-only fed male mice were measured after 1, 2, 3, 4, 5, 6, 7 weeks of feeding. (1w, 2w, 3w, 4w: CDAHFD n=5, egg n=5 per time point; 5w: CDAHFD n=4, egg n=4; 6w, 7w: CDAHFD n=4, egg n=3). (c) Photographs were taken before and after opening the abdomen. MCD on the sheets indicate modified MCD (CDAHFD). (d) Organ weights of CDAHFD fed or egg-only fed mice were measured. (e) Liver tissues removed at each time point were fixed in 4% paraformaldehyde and blocked in paraffin. Paraffin-embedded 4-μm sections were stained with hematoxylin and eosin and were observed on an Olympus light microscope and pictures were taken. (f) Levels of the liver enzyme markers ALT and AST were measured at 7 weeks. Data shown are the mean ratios ± standard errors. Statistical comparisons were performed using a Student T-test: -values<0.05 were considered statistically significant. * p-values<0.05, ** p-values<0.01.

**Figure 2.**
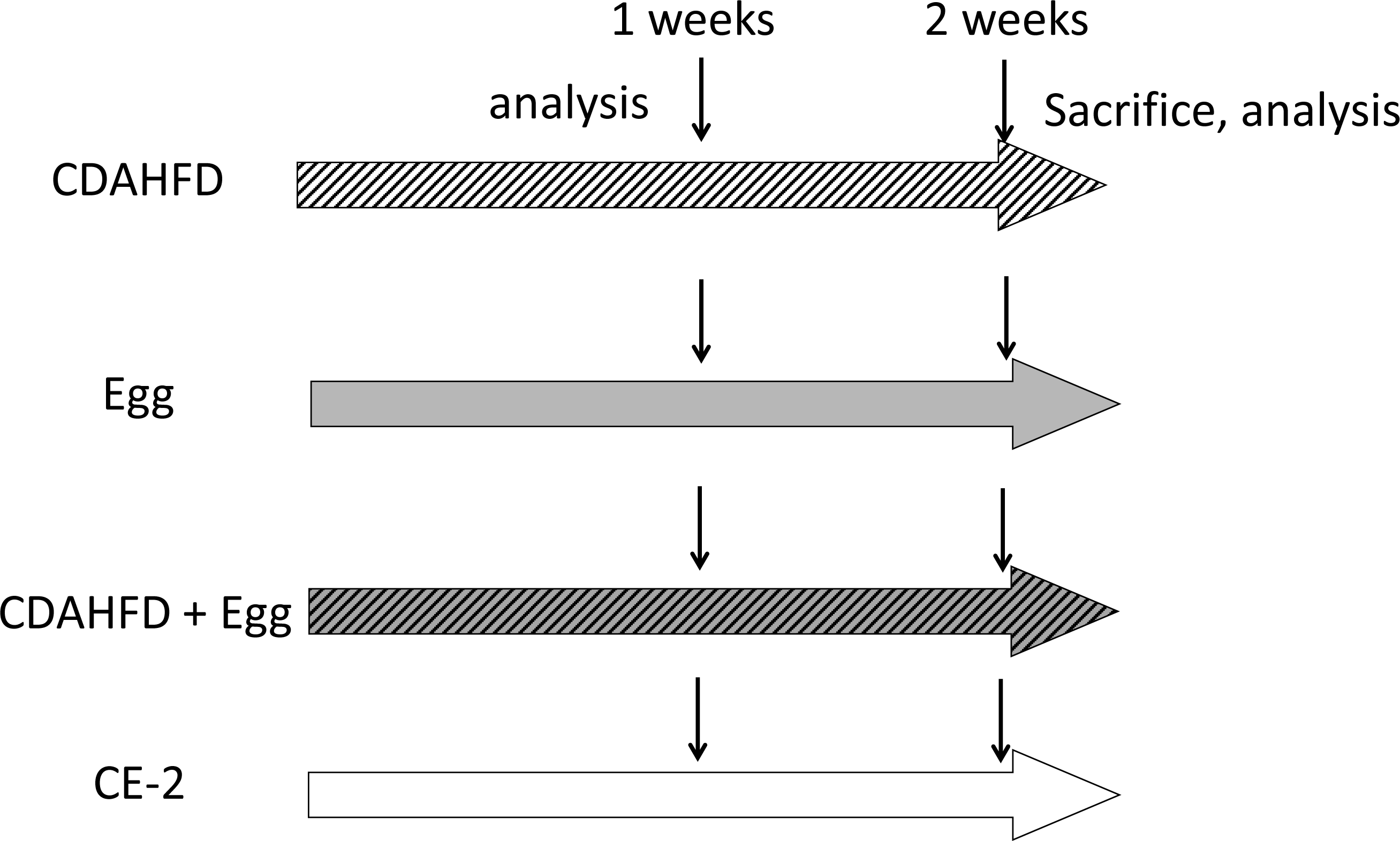
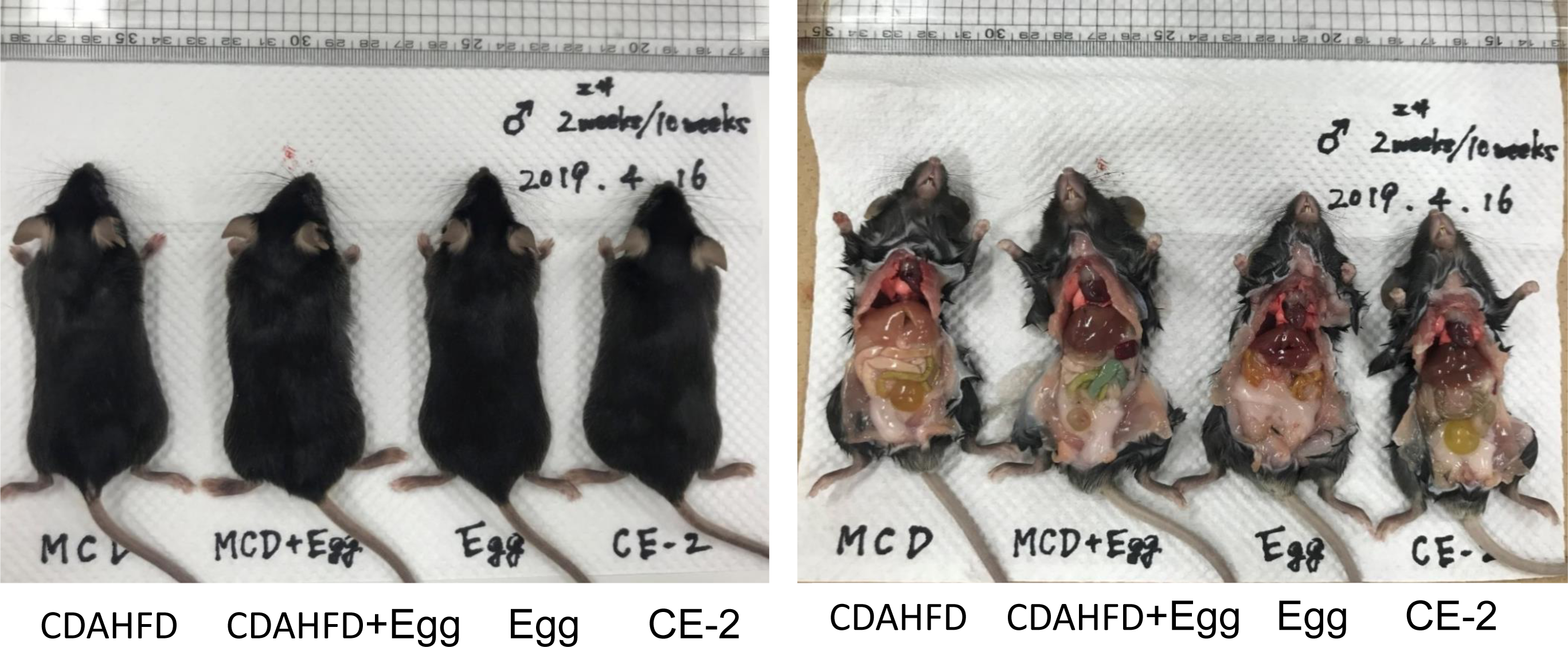
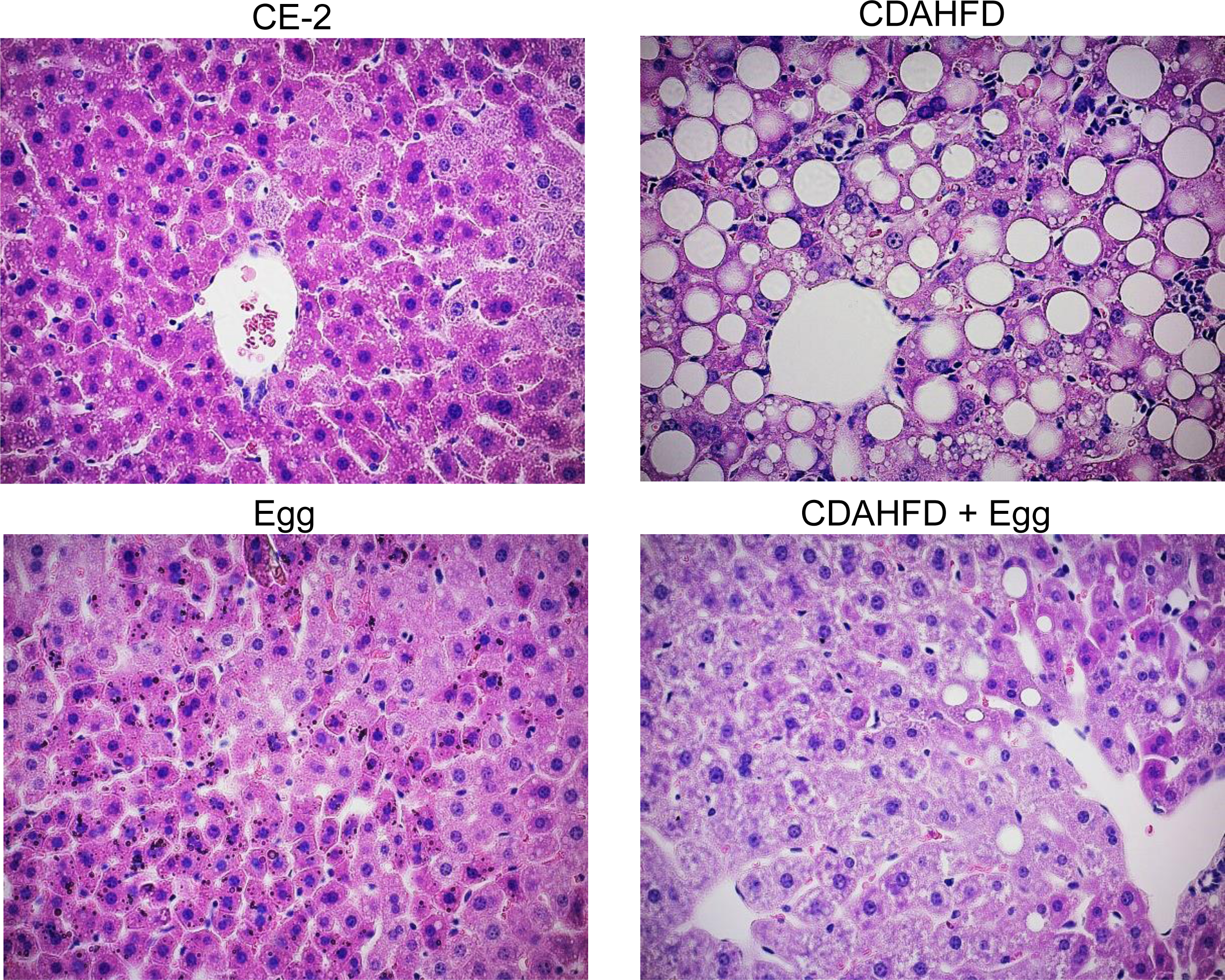
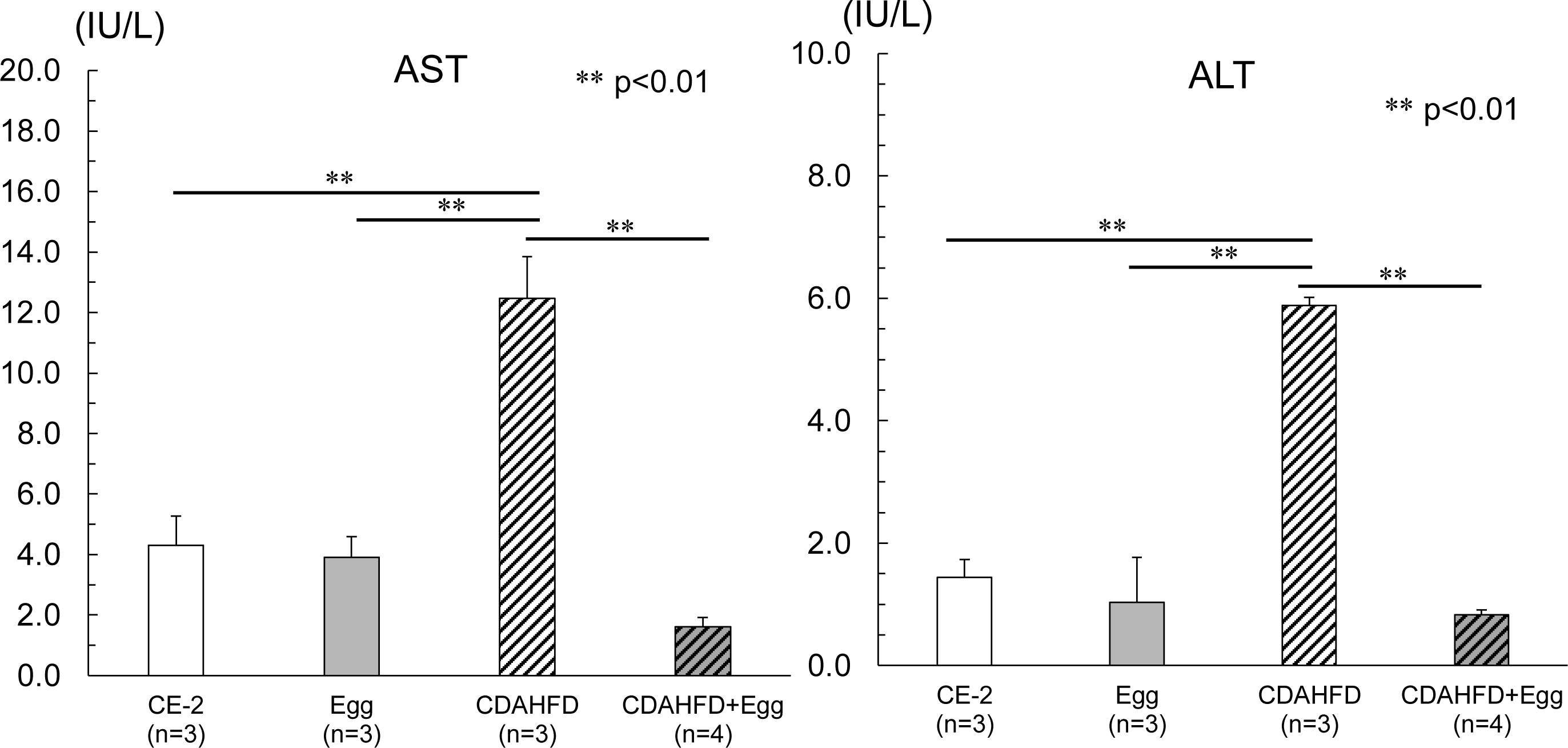
The addition of eggs to a CDAHFD prevents the induction of NAFLD. (a) Eight-week-old male mice were fed one of the following diets: CDAHFD, egg only CDAHFD and eggs or CE-2 ad libitum for 2 weeks. (b) Photographs were taken before and after opening the abdomen. MCD on the sheet indicates modified MCD (CDAHFD). (c) Liver tissues removed at each time point were fixed in 4% paraformaldehyde and blocked in paraffin. Paraffin-embedded 4-μm sections were stained with hematoxylin and eosin and were observed on an Olympus light microscope and pictures were taken. (d) Levels of the liver enzyme markers ALT and AST were measured at 2 weeks. Statistical comparisons were performed using a one-way analysis of variance (ANOVA) and Tukey-Kramer tests. P-values<0.05 were considered statistically significant. * p-values<0.05 ** p-values<0.01.

### The addition of eggs to a CDAHFD prevents the induction of NAFLD

Next we fed mice with a CDAHFD, egg-only or CE-2 diet for 2 weeks. In one group mice were fed both eggs and CDAHFDs (Fig.2a). In one week weight of the mice fed both eggs and CDAHFDs were higher than the other group at week one (data not shown). At two weeks all mice were appeared healthy. By opening the abdomen, we found that the livers of the mice fed the CDAHFD looked whitish and bigger than those of the other groups (Fig.2b). Liver weight in the CDAHFD fed mice was higher than that in the other group (Table 1). Liver histology showed that by eating a CDAHFD for 4 weeks, liver cells contained big and small vacuoles; in contrast the livers of egg-only fed mice and CE-2 fed mice had an almost normal appearance. Interestingly, adding eggs to the CDAHFD suppressed vacuole formation (Fig.2c). Serum ALT and AST in the mice fed an egg-only diet were normal, similar to values in the CE-2 mice. In contrast serum ALT and AST in the mice fed a CDAHFD were abnormally high indicative of hepatocytes death (Fig.1d).

**Table 1.**
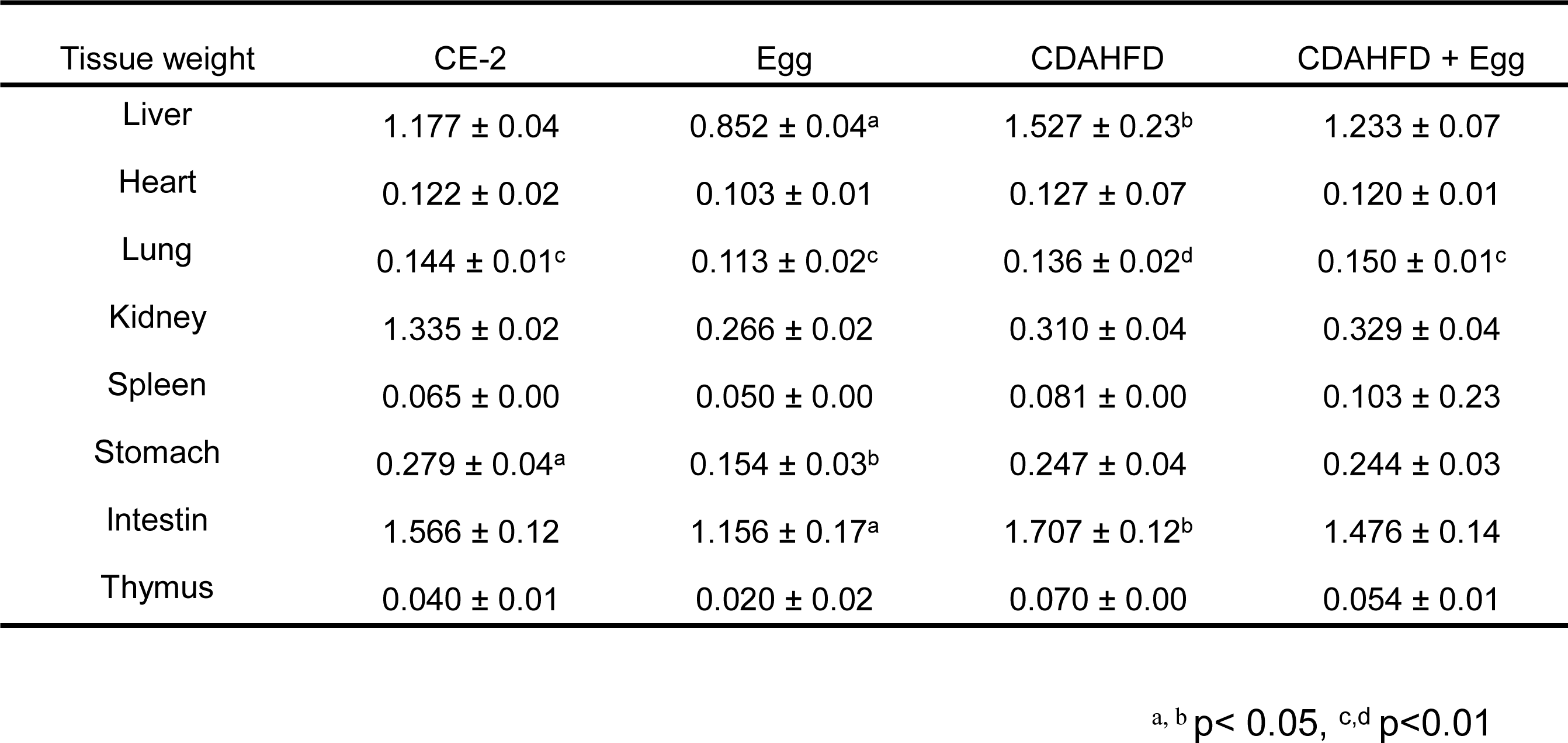
Eight-week-old male mice were fed one of the following diets: CDAHFD, egg only CDAHFD and eggs or CE-2 ad libitum for 2 weeks. Organ weights from each group of the mice were measured. Data are expressed as the means □±□ standard deviation. Statistical comparisons were performed using a one-way analysis of variance (ANOVA) and Tukey-Kramer tests. P-values<0.05 were considered statistically significant. p-values<0.05 **(a,b) p-values<0.01(c, d).

### Two weeks of an egg-only diet reversed the fatty liver caused by two weeks of a CDAHFD

Because we found that the CDAHFD induced a high grade NAFLD within only 2 weeks and considering the possible effect of eggs in suppressing fat accumulation in hepatocytes, we next examined whether egg-only diet could reverse the NAFLD caused by a CDAHFD. Eight-weeks-old male mice were first fed a lipid-rich diet for 2 weeks, then they were fed eggs only for another 2 weeks (Fig. 3a). We found that weights of the mice fed eggs only diet were higher than the other group at 3 and 4 weeks (Fig.3b). We found that the mice fed an egg-only diet for 2 weeks after 2 weeks of a CDAHFD recovered to almost normal liver conditioned based on results from liver appearance assessment, liver histology and serum ALT or AST obtained blood samples (Fig. 3c, d, e).

**Figure 3.**
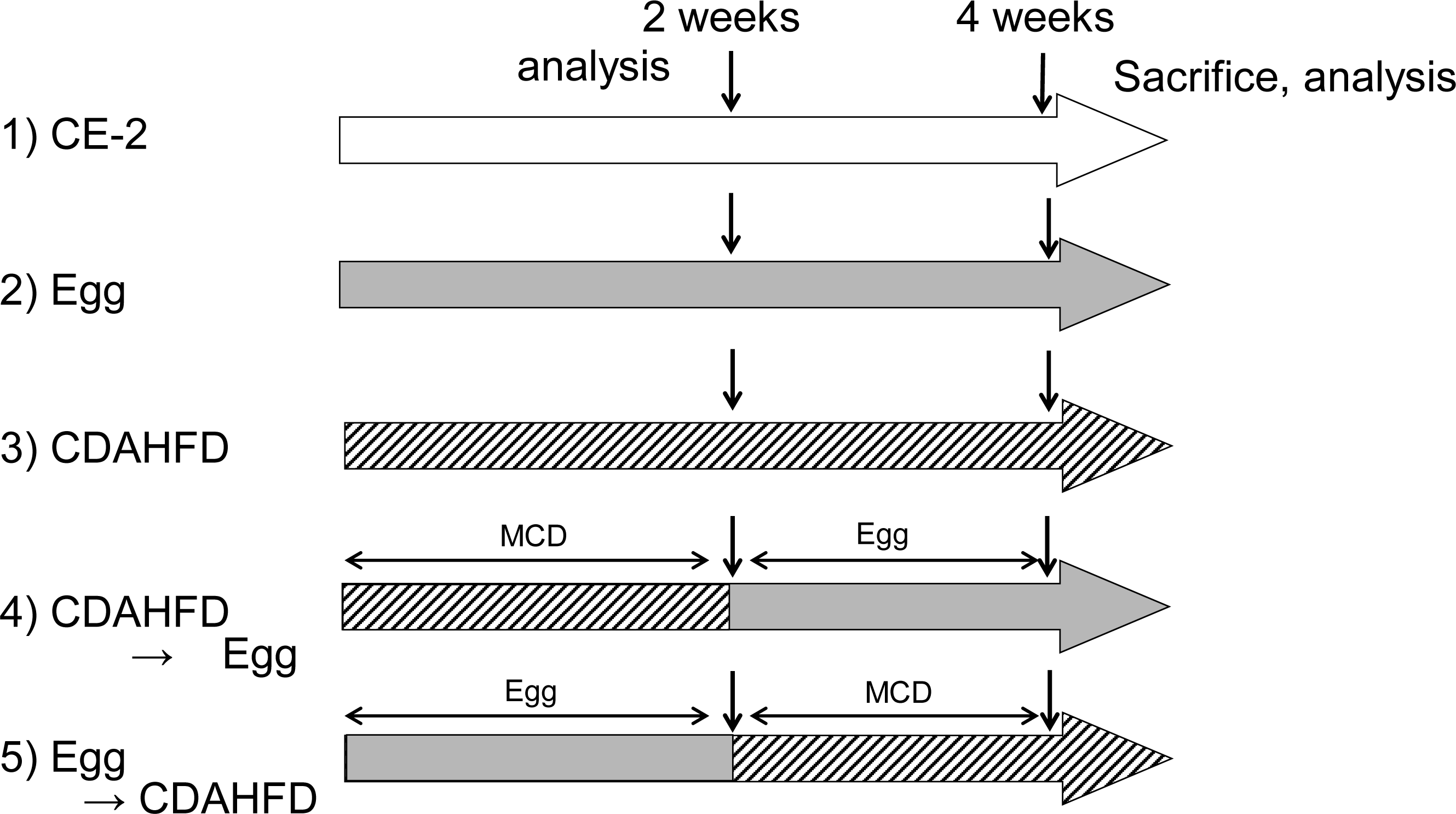
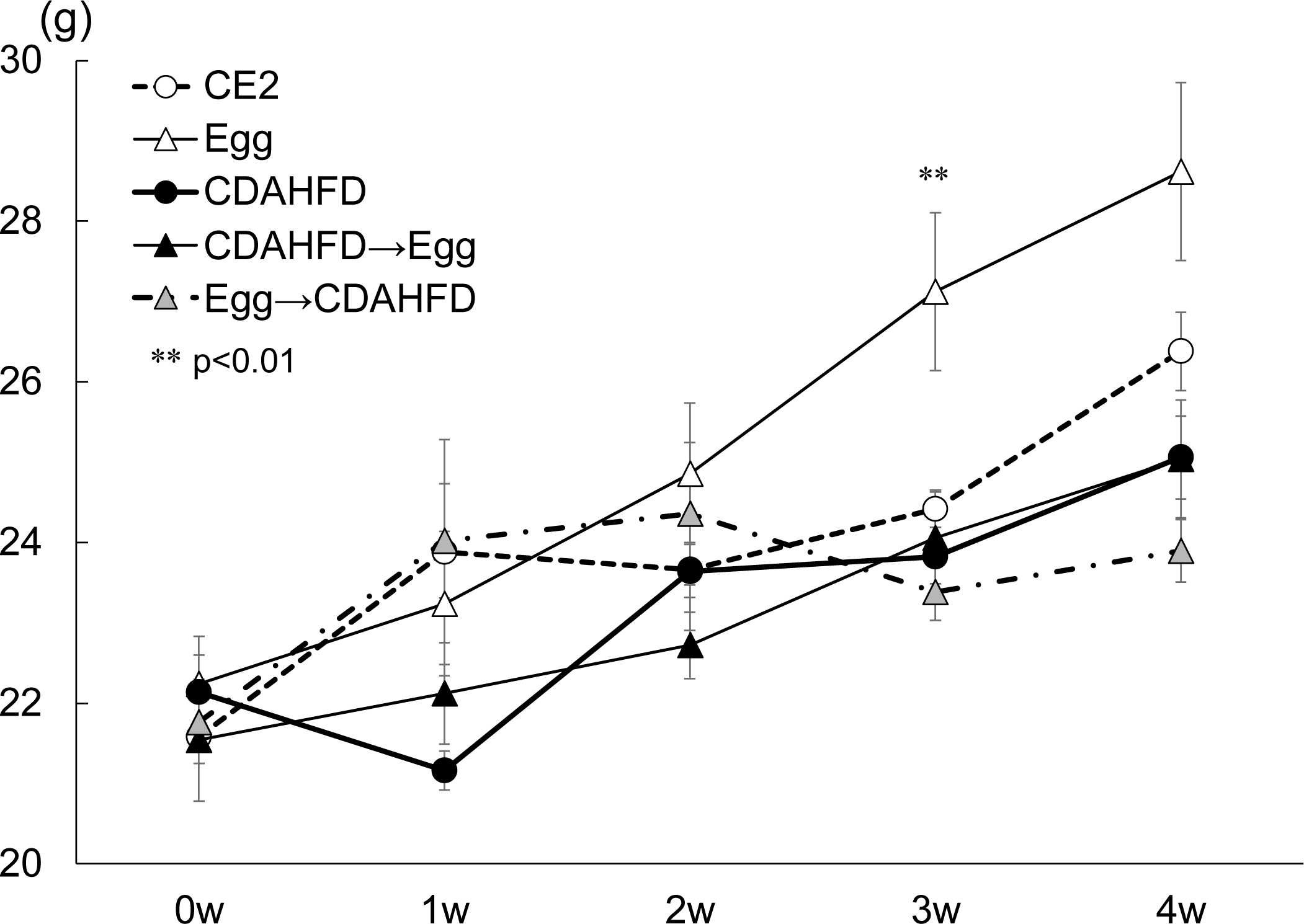
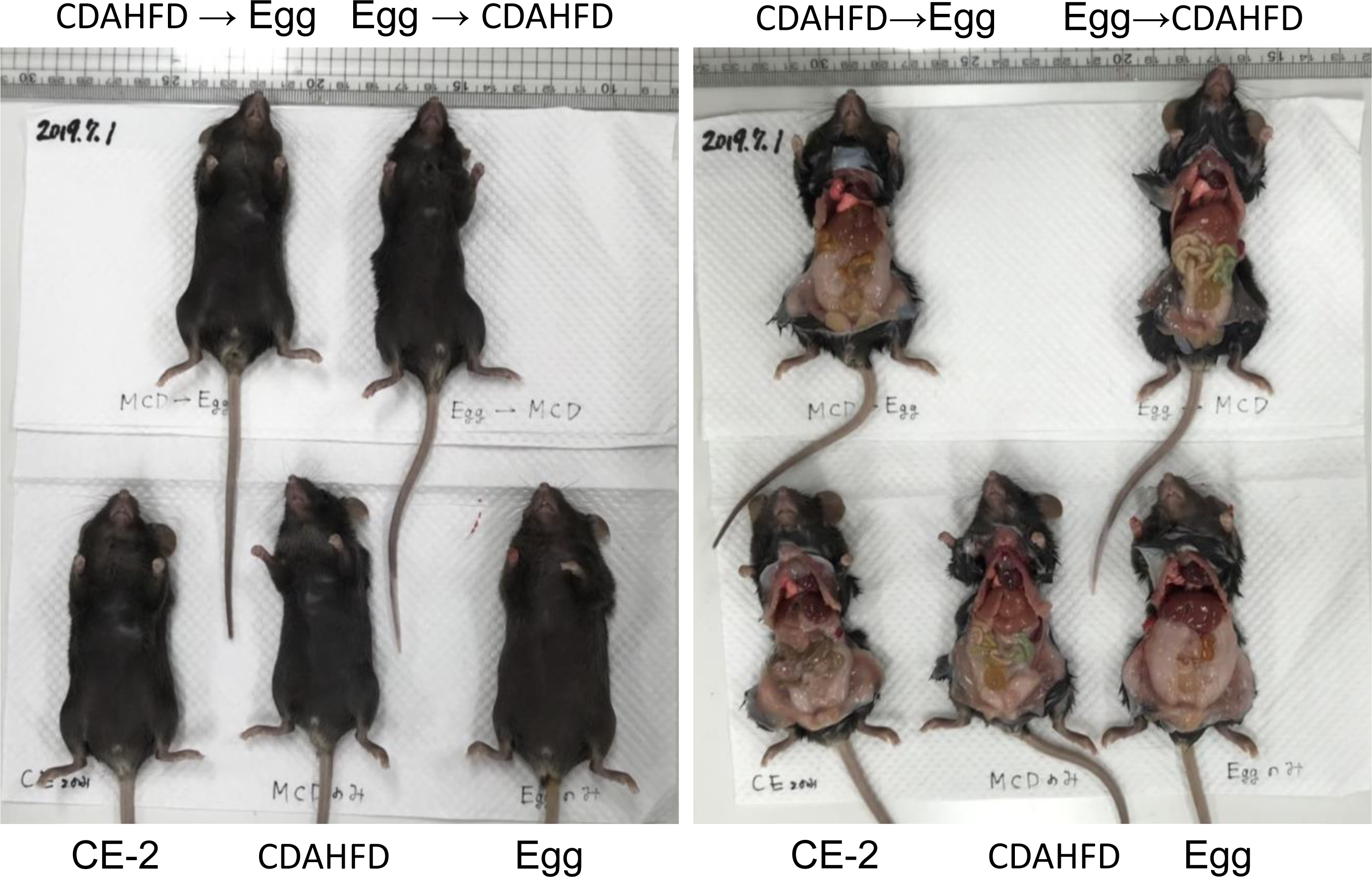
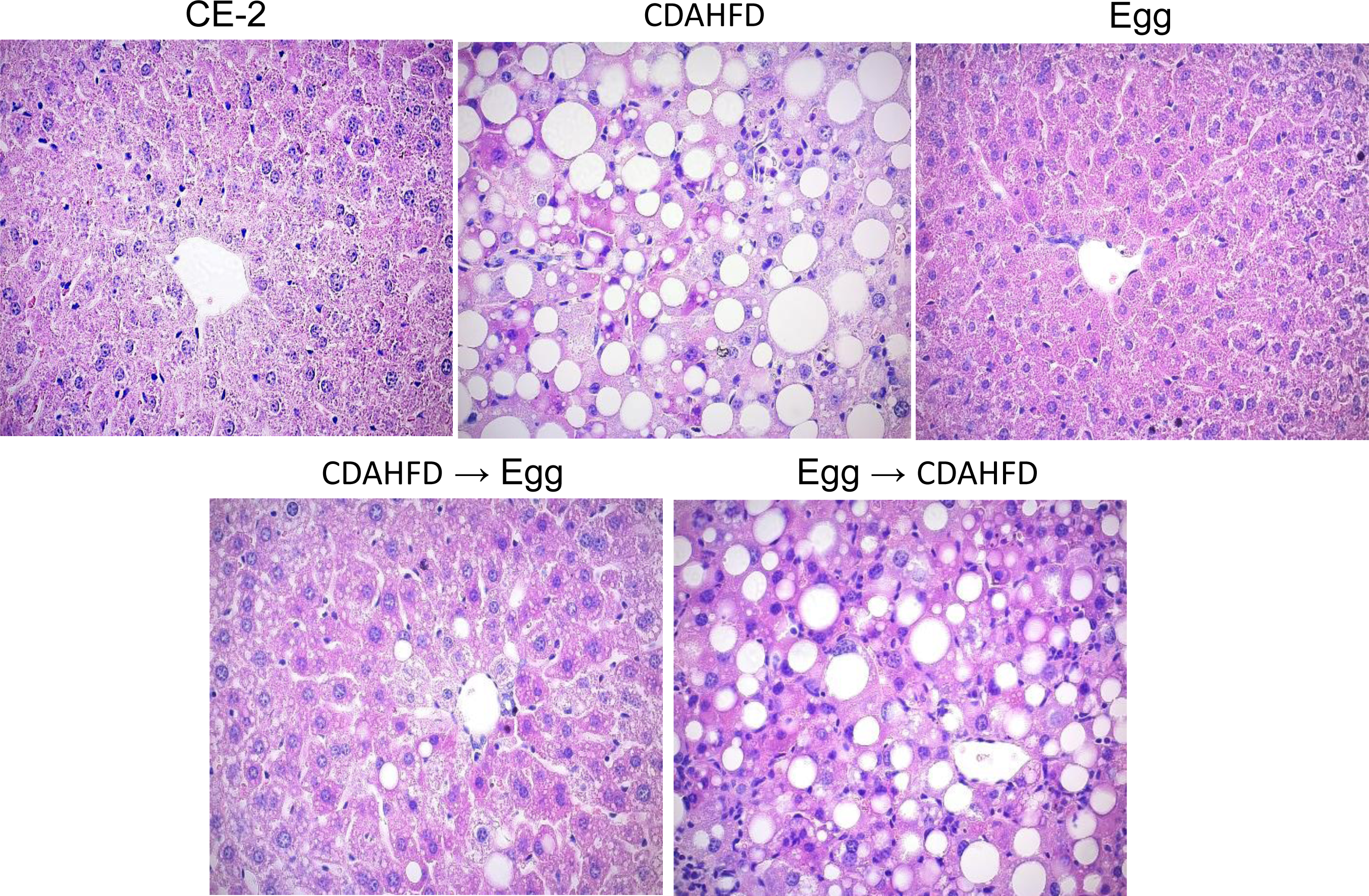
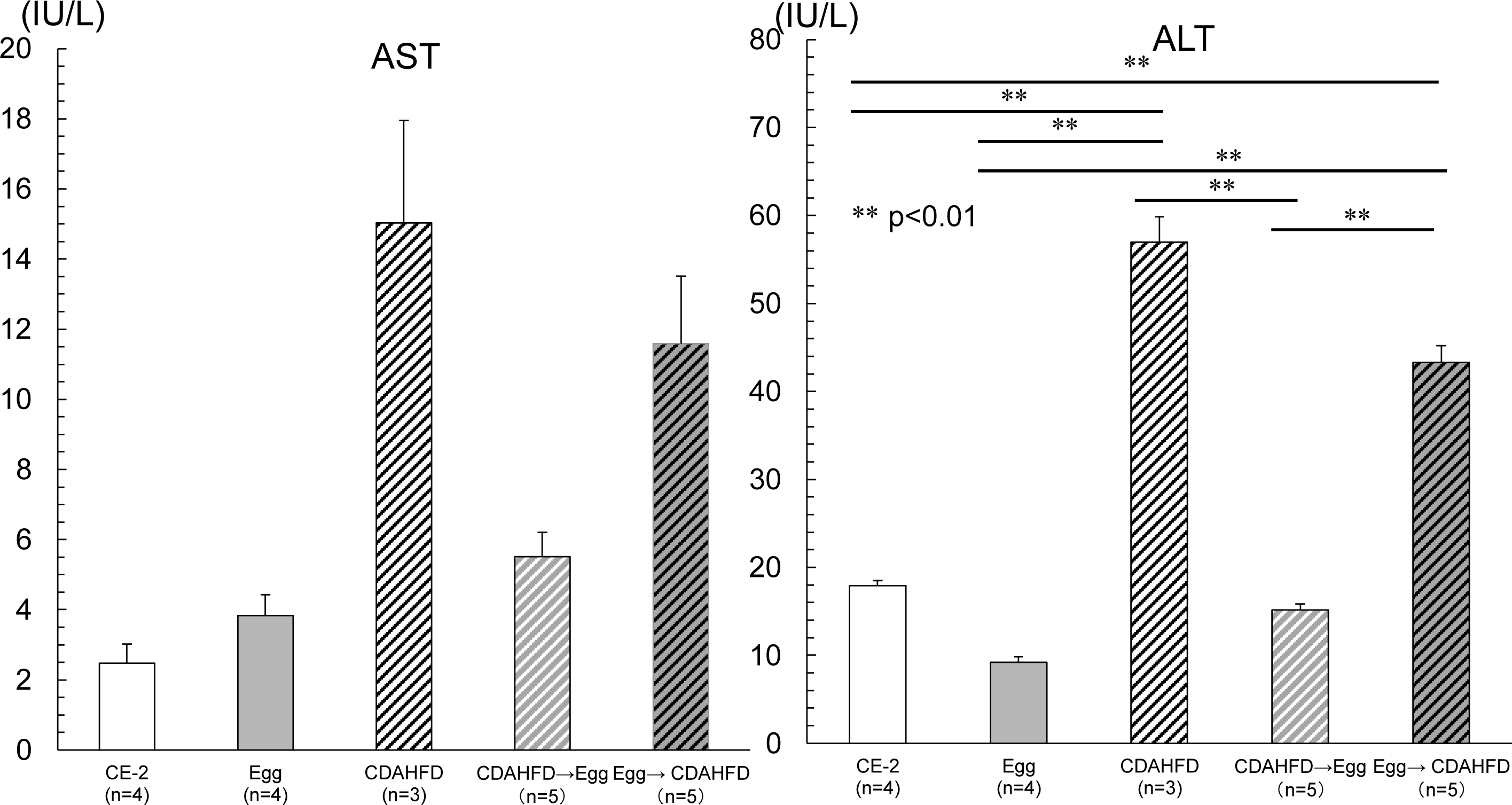
Eggs suppresses NAFLD after two weeks on CDAHFD diet. (a) Eight-week-old male mice were fed on the following diets: 1) CE-2, 2) CDAHFD, 3) eggs only, 4) CDAHFD for 2 weeks then switched to an egg-only diet for 2 weeks or 5) an egg-only diet for 2 weeks then switched to a CDAHFD for 2 weeks. (b) Body weights of each group were measured after 1, 2, 3, 4 weeks of feeding. Data shown are the mean ratios ± standard deviations (n=5). (c) Photographs were taken before and after opening the abdomen. MCD on the sheets indicate modified MCD (CDAHFD). (d) Liver tissues removed at each time point were fixed in 4% paraformaldehyde and blocked in paraffin. Paraffin-embedded 4-μm sections were stained with hematoxylin and eosin and were observed on an Olympus light microscope and pictures were taken. (e) Levels of the liver enzyme markers ALT and AST were measured at 2 weeks. Statistical comparisons were performed using a one-way analysis of variance (ANOVA) and Tukey-Kramer tests. P-values<0.05 were considered statistically significant. * p-values<0.05 ** p-values<0.01.

**Figure 4.**
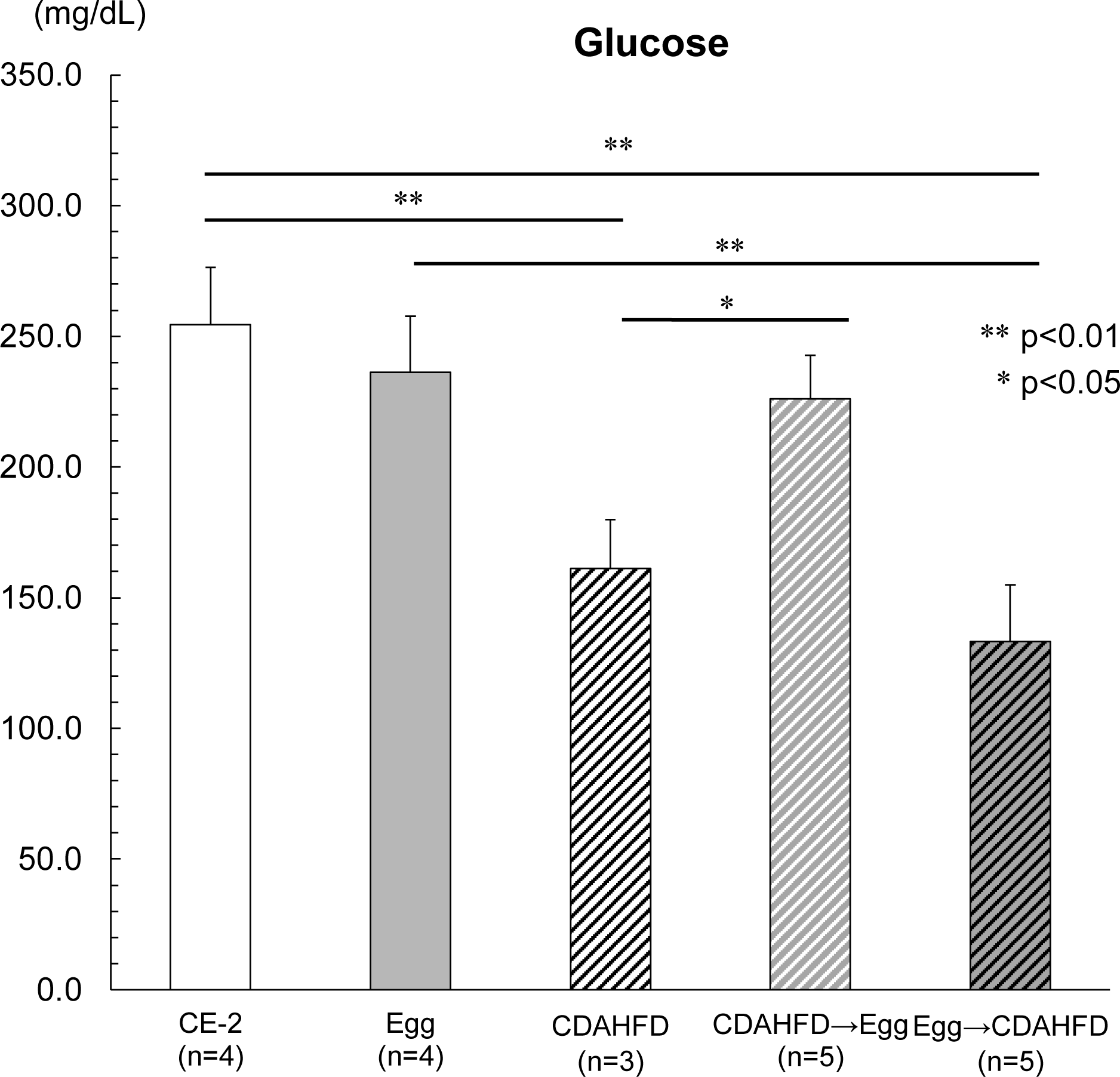
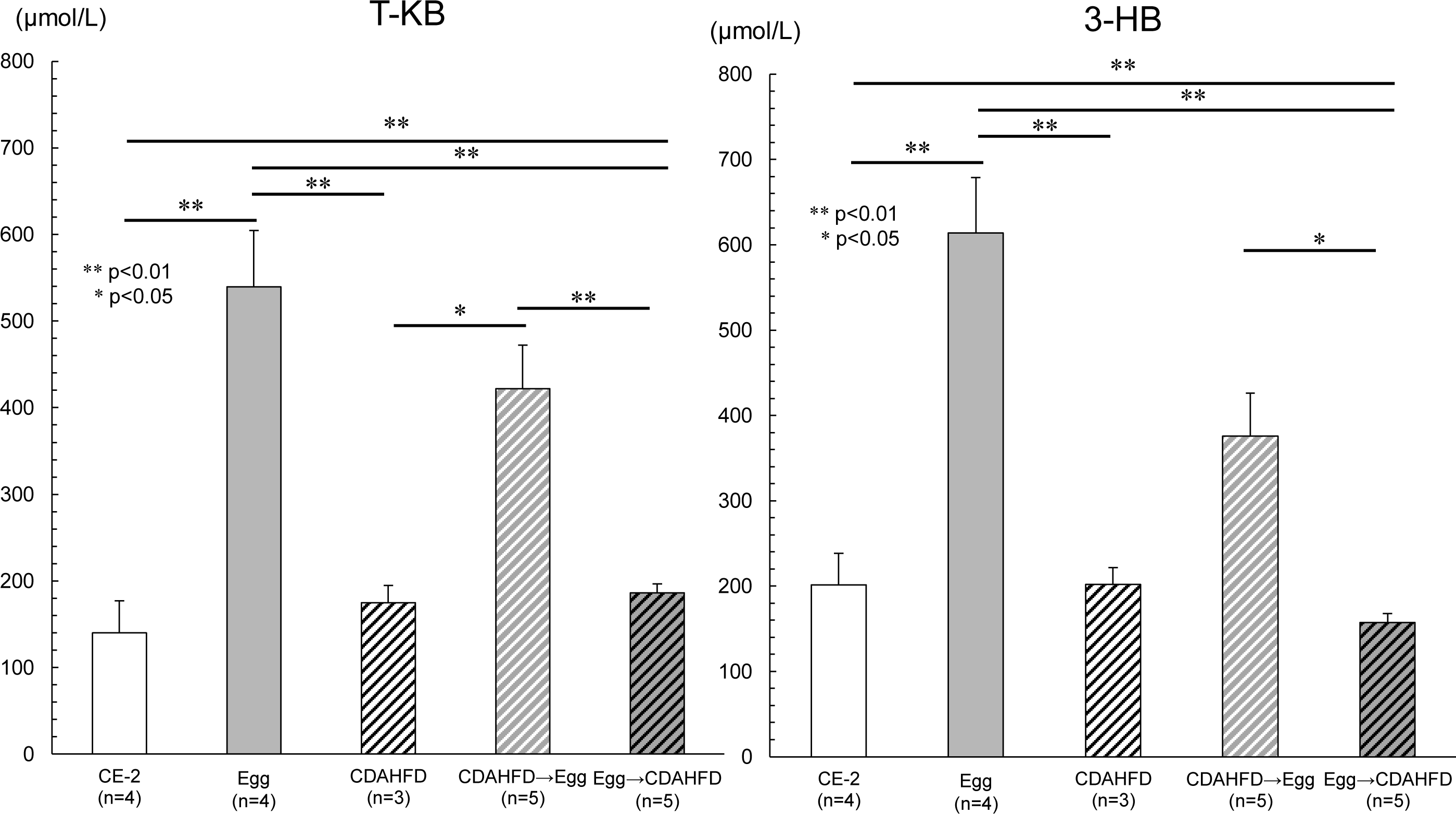
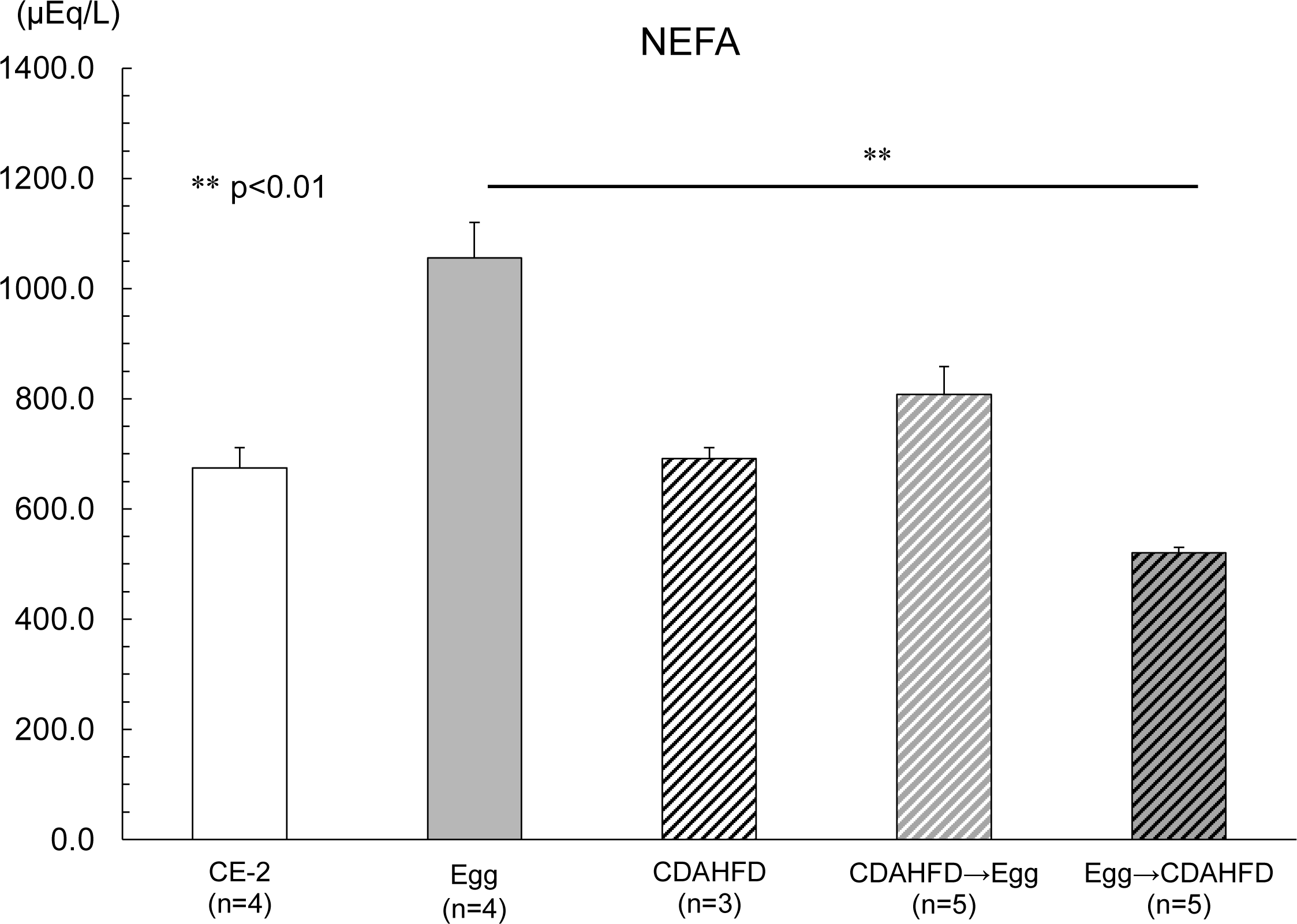
Comparison of serum glucose, ketone bodies and fatty acids between egg-only mice and CDAHFD mice. Serum from the mice in Fig.3 were taken in the morning after 2 weeks of each diet. (a) Levels of the serum of glucose. (b) Levels of the serum of T-KB and 3-HB. (c) Levels of the serum of free fatty acids. Statistical comparisons were performed using a one-way analysis of variance (ANOVA) and Tukey-Kramer tests. P-values<0.05 were considered statistically significant. * p-values<0.05 ** p-values<0.01.

### Comparison of serum glucose, ketone bodies and fatty acids between egg-only mice and CDAHFD mice

The serum glucose levels of the egg-only mice were almost the same as those of the CE-2 mice, whereas serum glucose of the CDAHFD mice was lower than that of the egg-only or CE-2 mice. The lower serum glucose levels in CDAHFD mice were reversed by switching to an egg-only diet (Fig.4a).

Both T-KB and 3-HB were much higher in the serum of egg-only mice than those in CE-2 or CDAHFD mice. Normal ketone bodies in the serum of CE-2 or CDAHFD mice increased after 2 weeks on an egg-only diet (Fig.4b). Serum fatty acids were also higher in the serum of egg-only mice than those in CE-2 or CDAHFD mice. The higher serum fatty acids levels in egg-only mice were decreased by switching to an CDAHFD diet (Fig.4c).

## Discussion

Here we have shown that an egg-only diet, which contains 60% fat and less than 5% of carbohydrates, does not induce NAFLD after 7 weeks. In contrast, a CDAHFD, which contains 60 % fat, induces NAFLD after only two weeks similar to a previous report (16). An HFD, which contains 60% fat has been used to induce NAFLD, although it takes a long time to develop fatty liver conditions (17).

There are several possible explanations as to why the same amount of lipid from an egg-only diet compared to a CDAHFD leads to different results. First, egg contains a lot of choline, whereas the CDAHFD specifically lacks choline. A high accumulation of triglycerides in the liver leads to fatty liver and triglycerides in the liver are derived from hepatic *de novo* lipogenesis (DNL) and high-fat diet (18).

Triglycerides in the liver are introduced into very low-density lipoproteins (VLDL) and secreted into circulation. If production and entrance of triglycerides in the liver exceed the secretion of triglycerides, triglycerides accumulate in the hepatocytes as fat droplets. VLDL is a lipoprotein having a single-layer phospholipid in outer shell, and which contains triglycerides and cholesterol-ester internally. Apolipoproteins are situated in the outer shell. Apolipoprotein B100, which is an important apolipoproteins in VLDL, may induce the incorporation of triglycerides into lipoprotein. Choline is essential for biosynthesis of phosphatidylcholine (PC), and is needed for the export of triglycerides (TG) out of hepatocytes via VLDL (19, 20, 21, 22). In humans, dietary choline restriction for three weeks induces liver damage (elevation of ALT) (23). Because choline (2-hydroxyethyl-trimethyl-ammonium salt; molecular weight of 104 g/mol) has several important functions in the human body and that in vivo biosynthesis of choline is not sufficient for daily life, choline needs to be obtained by diet (24). In dietary food, choline exists in water soluble form (free choline, phosphocholine, and glycerophosphocholine) and lipid-soluble forms (Phosphatidylcholine and sphingomyelin) (24). A boiled egg contains 225.7mg total choline mainly in the form of phosphatidylcholine (210.0mg in 100g of weight) (24,25). Several reports of dietary choline intake have been published. Mean dietary choline intake in healthy adult is around 300 to 500 mg /day; for instance 372 ± 287mg /day (Canadian men), 292 ± 213 (Canadian women), 445–513 (Japanese men), and 388–442 (Japanese women), which corresponds to about two boiled eggs (26,27). Our results presented here that high-fat containing chicken eggs did not induce fatty liver may be attributed to the high amount of choline in chicken eggs. Because the mice fed egg only consumed a very small amount of glucose, any DNL from glucose in the liver is trivial. However, these mice consumed a lot of triglycerides, which accumulated in the fat tissue, some of which entered the liver. The phosphatidylcholine (PC) in the eggs may encapusulate the triglycerides as a lipoprotein VLDL and the export them into circulation.

## Acknowledgments

This research was supported by 16H05280 and 19K11687 of the Ministry of Education, Science, Technology, Sports and Culture, Japan.

